# First Report of Nipah virus shed in urine by fruit bats *(Pteropus medius)*, Sri Lanka

**DOI:** 10.1101/2024.07.31.605971

**Authors:** Claudia Kohl, Sahan Siriwardana, Therese Muzeniek, Thejanee Perera, Dilara Bas, Mizgin Öruc, Annika Brinkmann, Beate Becker-Ziaja, Franziska Schwarz, Hamsananthy Jeevatharan, Jagathpriya Weerasena, Shiroma Handunnetti, Inoka C. Perera, Gayani Premawansa, Sunil Premawansa, Wipula Yapa, Andreas Nitsche

## Abstract

Nipah virus is an emerging pathogen with high public health relevance, causing outbreaks in South- and Southeast Asia with high case fatality rates. The natural reservoir host are fruit bats (*Pteropus spp*). In this study, we confirmed the presence of Nipah virus for the first time in Sri Lanka and are proving the genetic relatedness to Nipah virus strains from southern India, causing outbreaks in 2007, 2018, 2019 and 2023. It is noteworthy that the shedding of the Nipah Virus may have a temporal variation that correlates with the breeding season of *Pteropus medius*. Furthermore, preventive measures and possible mechanisms influencing viral shedding are discussed.

## Introduction

Anthropogenic climate change affects ecological systems globally and hence spatial biodiversity (1). From the infection biological point of view, affected biodiversity results in changed availabilities of susceptible vectors and hosts. Considering zoonotic Emerging Infectious Diseases (EID) in this context, involves many distinct factors. This is what the ‘One Health’ approach describes; the connectivity between human-, animal- and planetary health. There are several prominent examples of EIDs, enlightening this interdependence (e.g. Mpox, Hendra virus, Marburg virus, SARS Coronavirus and Nipah virus) (2-7).

Nipah virus (NiV, *Henipavirus nipahense*) and Hendra virus (HeV, *Henipavirus hendraense*) are species within the genus Henipavirus of the family *Paramyxoviridae* (8). Several Paramyxoviruses (i.e. Measles virus and RSV) are causing rather serious respiratory illnesses, and are often highly transmissible via the air-borne route. As an example, Measles virus has an R_0_ value of 12-18 and is as contagious that >90% of exposed non-immunized persons will develop symptomatic disease (9).

NiV has also proven its ability to cause even more severe outbreaks in humans with a broad range of clinical symptoms, ranging from subclinical infection to severe encephalitis, respiratory diseases and death (10). The case-fatality-rate (CFR) of NiV encephalitis is described as 61% (96% CI 45.7-75.4%)(10, 11). NiV first emerged in 1998 in Malaysia and Singapore (3). Bats (Flying foxes; *Pteropus spp*.) are the known reservoir hosts of NiV (12). Anthropogenic factors, such as land-use changes and deforestation may have forced remote living colonies of bats to relocate near human dwellings. Pigs, living in adjacent colonies were exposed to bat excretions and became intermediate hosts. Subsequent outbreaks were reported from Bangladesh, the Philippines and India resulting in more than 643 laboratory confirmed infections and 380 fatalities (10). Transmission from bat to human occurs via urine through different routes. Either by consumption of contaminated raw palm sap products, by direct contact with infected intermediate hosts (e.g. swine) or possibly by direct contact with bat urine (12). Though only five countries have reported NiV outbreaks so far, several more are on potential risk due to the distribution range of the reservoir host species (i.e. Bhutan, Indonesia, Maldives, Myanmar, Nepal, Sri Lanka, Thailand, Timor-Leste)(13). In 2023 several cases of Nipah virus were reported from Kozhikode (Kerala district in India) and the districts Dhaka, Rajbari and Shariatpur (Bangladesh). The shortest spatial distance between Sri Lanka (Mannar island) and India (Natarajapuram) is less than 55 km and *Pteropus spp*. are reported to migrate more than 450 km with ease (14). The importance of Nipah virus has been underlined by the recently published WHO South-East Asia Regional Strategy for the prevention and control of Nipah virus infection, 2023-2030, that is supposed to provide guidance to prevent severe illness and death from Nipah virus. In this study, urine excreted by several colonies of *Pteropus medius* in Sri Lanka was monitored for the presence of viruses using molecular techniques. A special focus was set on NiV, to investigate if NiV is present in Sri Lanka and to be able to take measures that prevent further spread.

## Methods

Between March 2018 and June 2019 bat colonies of *Pteropus medius* were monitored for the shedding of pathogens in Colombo, Mannar, Anuradhapura and Badulla, Sri Lanka. Clean sampling sheets were laid below roosting trees in the early morning and dripped urine was transferred drop by drop to sterile microtiter plates in the morning using disposable pipettes. All work was conducted while wearing appropriate Personal Protective Equipment (PPE) as described before (12).

A total of 2,218 urine samples were collected and combined into 32 pools with up to 8 urine specimens each. RNA was extracted from the individual pools using the QIAamp Viral RNA Mini Kit (QIAGEN, Hilden, Germany). Nipah virus screening was performed using real-time PCR targeting the phosphoprotein gene (P-gene), as described before (15). As an additional confirmation a real-time PCR assay, targeting the N-protein of the Nipah virus genome was developed (primers, probes and cycling conditions are described in table 1).

**Table 1.**
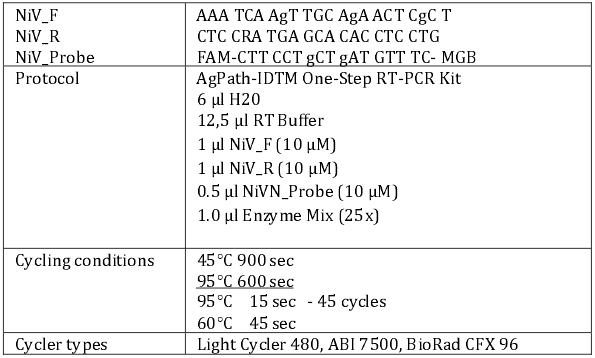
Primers and cycling conditions for the newly developed specific NiV qPCR assay.

The NiV positive pools S3_P02 and S18_P08 (2 of 32 pools) were subjected to Illumina HiSeq sequencing. Sequences were quality trimmed using Trimmomatic and mapped to NiV strain MCL-18-H-1088 (Acc. No. MHK523642, Kerala, India) as a reference strain. Additionally, an AmpliSeq primer panel (NiVliSeq), spanning the whole NiV genome, was developed using Primer3 (v 2.3.7). Primers are available on request.Corresponding PCR products were sequenced on the MinIon platform as described previously (16, 17). Draft genomes were submitted to NCBI Genbank (PP893186, PP893186, PP893186, PP893186) and aligned with NiV strains and type species of Paramyxovirus genomes, indicated by the ICTV and available in the NCBI GenBank database. Phylogenetic molecular clock reconstruction was calculated using MrBayes (MCMC approach), based on the GTR model (parameters: 1 Mio replicates, 4 chains, burn-in 10%) of the partial L sequence (7,977 nt; coding for the RdRP gene) of the subfamily Orthoparamyxovirinae, using Simian orthorubulavirus as an outgroup.

Ethics approval for this study was obtained from the Institute of Biology, Sri Lanka (WL/3/2/05/18) and necessary clearance was obtained from the Department of Wildlife Conservation, Sri Lanka.

## Results

Pool S3_P02 and pool S18_P08 were, both collected in Colombo and tested positive for NiV using the Henipavirus qPCR assay, targeting the P Gene (15). The NiV qPCR assay (targeting the N gene) further confirmed NiV in the same sample pools. All results are summarized in table 2.

**Table 2.** Overview on samples and sequencing results for NiV positive bat urine samples from Sri Lanka.

|  | S03-P2 | S18-08 |
| --- | --- | --- |
| Origin | *Pteropus medius*, Colombo | *Pteropus medius*, Colombo |
| Sampling date | June 2018 | March 2019 |
| PCR (CT value, mean duplicates) | Henipa PCR (33,8)<br>NiV in house (32,3) | Henipa PCR (35,6)<br>NiV in house (37,7) |
| Illumina Sequencing (Shotgun) |  |  |
| Trimmed reads | 5,094,516 | 5,633,085 |
| NiV reads | 128 reads | 20 reads |
| Longest contig | 481 nt (Acc. No XXXXX) | 255 nt (Acc. No XXXXX) |
| Highest percent ID nt | 98.96 MCL-18-H-1088<br>(Acc.No. MHK523642)<br>Human, outbreak 2018 Kerala | 99.61 (Acc.No. MN549404.1)<br>Bat India |
| NiVliSeq (Amplicon based approach) |  |  |
| Genome length and coverage | 93% query, 189-fold average<br>(> 10-fold) | 98% query, 8,783-fold average<br>(> 10-fold) |
| ID nt | 98.34% id nt (MH396625)<br>Human, outbreak 2018 Kerala | 98.43% id nt (MH396625)<br>Human, outbreak 2018 Kerala |
| Novel strain | Nipah virus strain C-18-B-0302 Sri Lanka<br>(Acc. No XXXXXXXX) | Nipah virus strain C-19-B-1808 Sri Lanka<br>(Acc. No XXXXXXXX) |

Metagenomic sequencing revealed corresponding NiV reads in these two sample pools, mapped all-over the NiV genome (table 2) and hence providing additional confirmation. The NiVliSeq AmpliSeq approach revealed 93 % of the NiV genome from pool S3_P02 and 98 % of NiV genome for pool S18-08, respectively (table 2). Comparison of the obtained NiV genomes with the NCBI Genbank database using BLASTn, revealed for both genomes the highest identity to strain MH396625 obtained from a human patient during the 2018 Outbreak in Kerala, India, with 98 % id nt and 98% id nt, respectively. The Sri Lankan NiV strains were named according to the systematic used for the Indian strains: NiV strain C-18-B-0302 Sri Lanka (Acc. No XXXXXXX) and NiV strain C-19-B-1808 Sri Lanka (Acc. No XXXXXXX); C for Colombo; 18 for 2018; 19 for 2019; B for bat; sample number.

Phylogenetic analysis clearly allocated the novel Nipah virus strains C-18-B-0302 Sri Lanka and C-19-B-1808 Sri Lanka to the species Nipah virus (*henipavirus nipahense*) within the genus Henipavirus (Figure 1). Both are clustering monophyletically with the strains sampled from humans during the outbreaks in Kerala region, India in 2007, 2018, 2019, 2021 and 2023.

**Figure 1:**
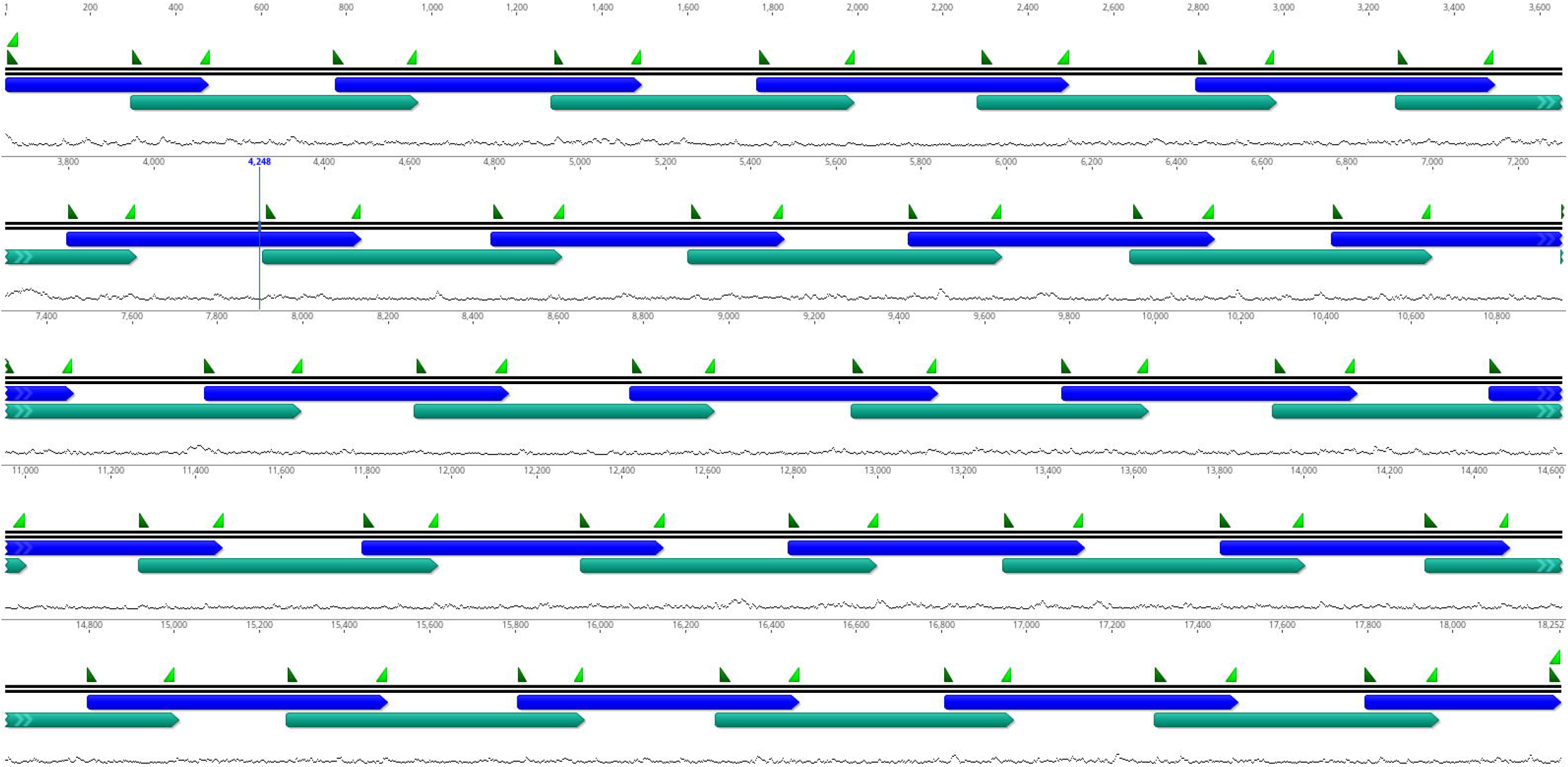
Phylogenetic reconstruction of Subfamily Orthoparamyxovirinae, using Simian orthorubulavirus as an outgroup. The alignment is based on the partial L gene (7,977 nt) coding for the RdRP. Calculation was done using MrBayes MCMC approach of molecular clock reconstruction, based on the GTR model, 1 Mio replicates, 4 chains, burn in 10%.

## Discussion

This study demonstrated the presence of public health-relevant NiV strains in Sri Lanka for the first time. The developed qPCR and NiVliSeq Sequencing approach validated the already obtained results and represent valuable tools, which will be used in further studies. The differences in genome coverage are seemingly not corresponding to the underlying CT values. This is due to the fact that the sample material for sample S03-P2 was limited and could not be used for further experiments. However, the very high genome coverage, even at the high CT levels of >35, reflecting lower viral loads, underlines the power of the methodological approach.

The identified strains have highest similarities to human-pathogenic strains causing recent outbreaks in India (2007, 2018, 2019 and 2023) and hence are of major importance for public health. However, NiV is still a rare disease and transmission rates are comparably low. Further, the detection of a certain virus in a bat population does not necessarily mean, that transmission occurs. Likely, diverse strains of Henipaviruses can be found in *Pteropus spp*. across the whole distribution range. Many factors are contributing to the successful shedding, transmission and infection. NiV is a perfect example, that zoonotic diseases have to be tackled via one health approach, taking all key factors into account: Animal health and wellbeing, climate change, human health and behavior.

The potential of zoonotic bat-to-human transmission could be minimized by preventive measures. Such measures could comprise education of healthcare workers to raise appropriate awareness for NiV, equipment and training for the early detection of NiV RNA in clinical specimens and preparation for clinical management of NiV-positive patients.

The initial prevention of any infection should be one of the major goals. Circular fencing of trees with roosting bats in public areas would likely prevent possible contact with bat urine. Bat colonies need to be protected for several reasons: among which the ecosystem services they provide including pollination. Moreover, *Pteropus spp*. are threatened by extinction (18). Additionally, it is important to realize, that emergence may be a result of disturbances of ecological systems and hence cannot be cured by further disturbance.

The World Health Organization (WHO) has just recently recognized the need for research and diagnosis of NiV within the distribution range of Pteropus spp. (22). Shedding of NiV likely also follows seasonal and temporal pulses like already described for Hendra virus (19-21). These pulses may either be explained by episodic shedding from persistently infected bats or transient shedding waves of bat-to-bat transmission between bat populations as a result of waning herd immunity (20). Immunity may be influenced by stress, such as weather events, reproduction cycles and anthropogenic pressure like land-use changes. Recently, a mechanism was identified which allows the retrospective explanation and prediction of Hendra virus shedding in Australia. Warming of the ocean (El Niño) influences and prevents the winter flowering of trees in remnant forests and hence causes nutritional stress for the bats (21). Eby et al. showed significant correlations between El Niño and Hendra virus shedding. Being a tropical country, Sri Lanka has complex and diverse ecosystems where their resilience to climate change may be better than other regions of the world. Hence, it is important to find out the reasons behind the temporal viral shedding observed in Sri Lanka. Similar mechanisms may be influencing the shedding of NiV and have to be understood from the One Health point of view, to prevent further spread.

---

Supplemental table 1

Position of primers used for the NiVliSeq Amplification Panel.

## Acknowledgments

We are grateful to the MF2 NGS core facility at the RKI for Illumina sequencing and Ursula Erikli for copy-editing. BMG funding, Department of Wildlife Conservation, Sri Lanka for necessary approvals and permits. This research was conducted according to the guidelines of the Fauna and Flora Protection Ordinance (FFPO) of Sri Lanka, under the permit No. WL/3/2/05/18 issued by the Department of Wildlife Conservation, Sri Lanka (10 January 2018). We also thank Colombo Municipal Council for granting permission and access to conduct the research in the public Park.

## Disclaimers

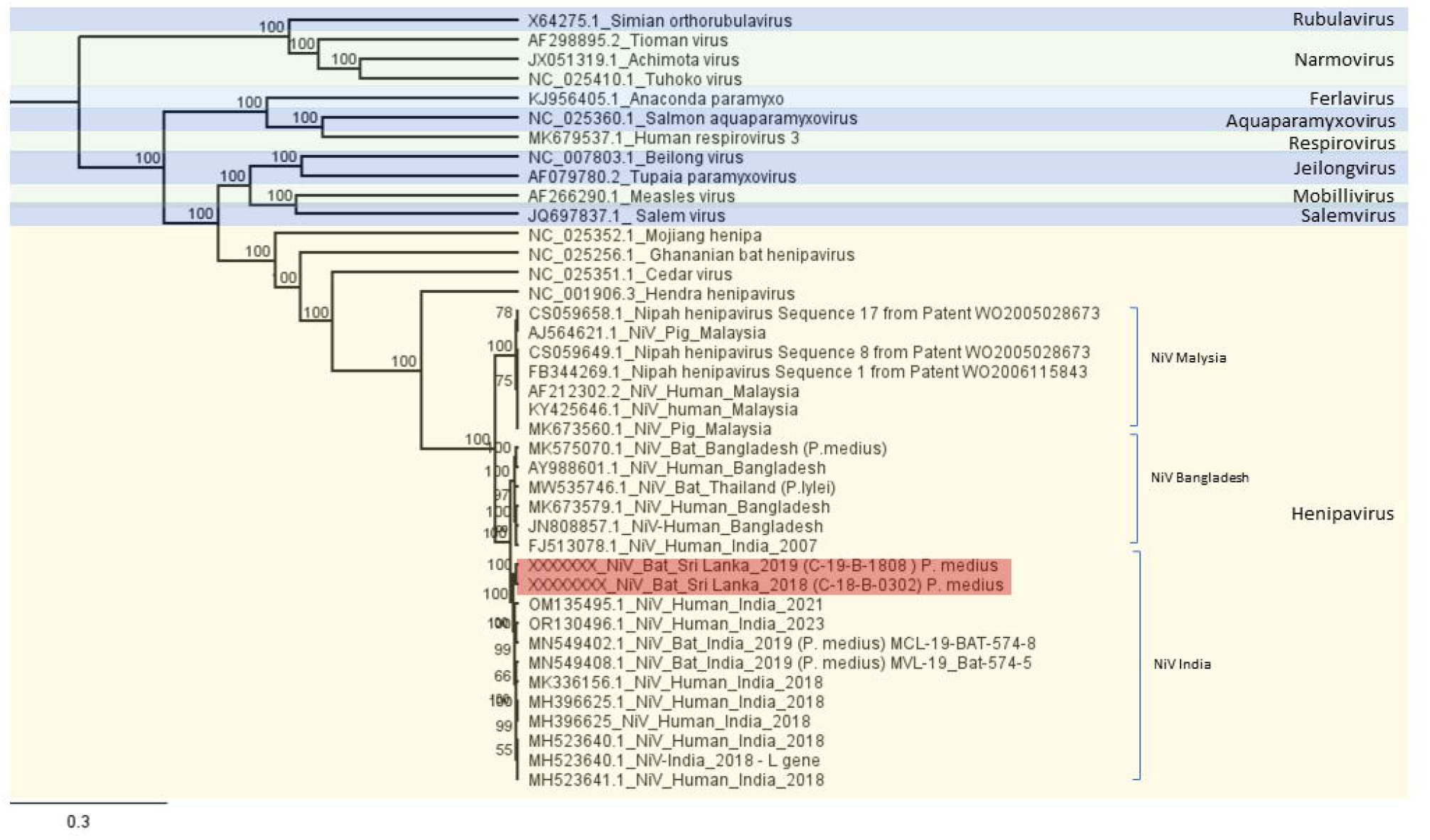

